# Antisense oligonucleotide therapy for *KCNT1* encephalopathy

**DOI:** 10.1101/2020.11.12.379164

**Authors:** Lisseth Estefania Burbano, Melody Li, Nikola Jancovski, Paymaan Jafar-Nejad, Kay Richards, Alicia Sedo, Armand Soriano, Ben Rollo, Linghan Jia, Elena Gazina, Sandra Piltz, Fatwa Adikusuma, Paul Q. Thomas, Frank Rigo, Christopher A. Reid, Snezana Maljevic, Steven Petrou

## Abstract

Developmental and epileptic encephalopathies (DEE) are characterized by pharmacoresistant seizures with concomitant intellectual disability. Epilepsy of infancy with migrating focal seizures (EIMFS) is one of the most severe of these syndromes. *De novo* mutations in ion channels, including gain-of-function variants in *KCNT1*, have been found to play a major role in the etiology of EIMFS. Here, we test a potential precision therapeutic approach in *KCNT1*-associated DEE using a gene silencing antisense oligonucleotide (ASO) approach. The homozygous p.P924L (L/L) mouse model recapitulates the frequent, debilitating seizures and developmental compromise that are seen in patients. After a single intracerebroventricular bolus injection of a *Kcnt1* gapmer ASO in symptomatic mice at postnatal day 40, seizure frequency was significantly reduced, behavioral abnormalities improved, and overall survival was extended compared to mice treated with a control ASO (non-hybridizing sequence). ASO administration at neonatal age was also well-tolerated and effective in controlling seizures and extending the lifespan of treated animals. The data presented here provides a proof of concept for ASO-based gene silencing as a promising therapeutic approach in *KCNT1*-associated epilepsies.

## INTRODUCTION

Epilepsy of infancy with migrating focal seizures (EIMFS) (previously known as malignant migrating partial seizures of infancy) is one of the most severe developmental and epileptic encephalopathy (DEE) syndromes (1). Pharmacoresistant, nearly continuous multifocal seizures begin to occur during the first six months of life and are accompanied by a marked developmental regression or stagnation (1, 2). In addition, major axial hypotonia, as well as pyramidal and extrapyramidal signs become more apparent with the progressive development of athetotic movements and other movement disorders (2–4). Many of these patients also display microcephaly and strabismus (2–4). The prognosis of this condition is very poor with most patients being non-verbal and non-ambulatory (2, 4, 5). Importantly, this syndrome is associated with a high mortality rate (ranging from 17% to 33%) (1, 2, 5), that can occur as a consequence of prolonged status epilepticus and respiratory failure (2).

Pathogenic gene variants have been identified in *KCNT1* and account for up to 50% of the etiology of EIMFS (1, 3, 6–10). *KCNT1* encodes the sodium activated potassium channel protein KNa1.1 which mediates an outward rectifying K^+^ current related to the slow hyperpolarization that follows repetitive action potential firing (11–16). Functional studies have shown that *KCNT1* pathogenic variants associated with epilepsy result in an overall gain of function effect on the channel activity, increasing the current up to 22-fold compared to the wild-type channel (6, 11, 17). For a subset of these pathogenic variants, changes in voltage dependence or Na^+^ sensitivity contribute to the increase in current (18, 19). Gain of function variants in *KCNT1* have also been found in patients with other epileptic syndromes including Autosomal Dominant Nocturnal Frontal Lobe Epilepsy, Ohtahara syndrome, and Lennox-Gastaut syndrome (3, 5, 7, 8, 20, 21). The antiarrhythmic drug quinidine has been shown to reduce the pathogenic currents produced by some of the mutant KCNT1 channels *in vitro* (11, 17, 22–25) but has had a limited clinical translation due to unwanted side effects (QTc prolongation and increased risk for arrhythmia) thereby resulting in a limited therapeutic window (25–37). To date, there is no effective therapy to treat *KCNT1* associated epilepsies.

RNA-targeted therapies have recently received significant attention for the recurrent examples of preclinical and clinical success, as well as fast-tracked development (38, 39). Some of these approaches are proving to be disease-modifying in the treatment of progressive neurological conditions (40, 41). Of the RNA-targeted therapies, which include antisense oligonucleotides (ASOs), short interfering RNA (siRNA), antagomirs, microRNA mimetics, and DNAzymes, ASOs have provided the most impactful clinical benefit (38, 41, 42). ASOs have emerged as a therapeutic option for orphan pediatric neurogenetic conditions including spinal muscular atrophy (43) and Duchenne’s muscular dystrophy (44). ASOs are synthetic, single-stranded nucleic acids typically of 10 to 30 nucleotides in length that bind to a specific complementary mRNA target sequence and modulate the expression of a specific gene at the RNA level (42). This control is achieved by different molecular mechanisms, from the steric block of ribosomal activity to regulation of RNA splicing (39, 42). ASOs that harness endogenouse RNase H1 mechanisms are commonly used to specifically reduce the expression of mRNAs. These ASOs are referred to as gapmer ASOs because they contain central block of deoxynucleotide monomers needed to induce the cleavage of target mRNA (39, 42). The major advantage of ASOs as a therapeutic approach is that the interaction with the target is considerably more specific than traditional small-molecule based therapeutics (42).

To investigate the therapeutic potential of reducing *KCNT1* expression we evaluated the efficacy of a *Kcnt1* gapmer ASO in a mouse model of *KCNT1*-DEE. Mice homozygous for the *Kcnt1* variant p.P905L (L/L) display spontaneous seizures, abundant interictal activity in the electrocorticogram (ECoG), behavioral abnormalities, and early death. After a single intracerebroventricular (i.c.v.) bolus injection of *Kcnt1* gapmer ASO at postnatal day 40, L/L mice showed a marked knockdown of *Kcnt1* mRNA, resulting in an almost complete abolition of seizures, prolonged survival, and improved performance in behavioral tests. A dose of gapmer ASO that produced >90% *Kcnt1* mRNA knockdown to mimic exaggerated pharmacology in wild-type mice was well tolerated suggesting minimal on-target liability. The preclinical evidence presented here provides proof of concept for ASO based gene silencing as a therapeutic approach in *KCNT1* gain of function epilepsies.

## RESULTS

### The *Kcnt1* L/L model of *KCNT1* encephalopathy

*Kcnt1* p.P924L heterozygous mice (L/+) showed no evident epileptic or behavioral phenotype (**Figures 1 and 2**) and no increased susceptibility to chemical or thermally induced seizures (**Extended data figure 1 a-d**). In contrast, homozygous mice (L/L) were smaller in size (**Figure 1 a-b**) and had a markedly reduced lifespan compared to their littermates (median survival of 43 days) (**Figure 1 c**). In addition, the yield of L/L mice, born from L/+ breeders, was not consistent with the predicted Mendelian ratio of 1:2:1 (+/+: L/+: L/L) with only 8.8% of pups born found to be L/L by age P8-12 (age at which mice are sampled for genotyping) (**Extended data Figure 1 e**). Spontaneous tonic-clonic seizures were observed as early as P18. Video monitoring and handling showed diverse seizure phenotypes, with milder seizures lasting approximately 1 to 2 minutes and involving clonic movements of the forelimbs, neck and head while in a seated posture. More severe seizures including generalized tonic-clonic episodes with Straub tail, wild running and jumping, loss of postural tone and tonic hindlimb extension. Isolated tonic seizures (pure postural tone loss) were also observed. Milder seizures, in general, were not followed by an evident postictal state, and the mice quickly recovered. Longer and generalized seizures were followed by a short period of complete immobility and rapid breathing with later episodes of reduced mobility lasting up to 10 minutes. Status epilepticus (a seizure lasting > 5 minutes) was also observed and in most cases resulted in death. The frequency of tonic-clonic seizures was variable between mice (**Figure 1 f**). ECoG recordings revealed the presence of frequent high amplitude interictal acute spikes in the L/L mice, (**Figure 1 d**) this signal was absent in L/+ and +/+ (**Figure 1e**) and did not correlate with changes in behavior. Convulsive seizures were associated with ictal ECoG signals characterized by clusters of high-amplitude sharp wave activity followed by electrical suppression at the end of the seizure (**Figure 1 d**).

**Figure 1.**
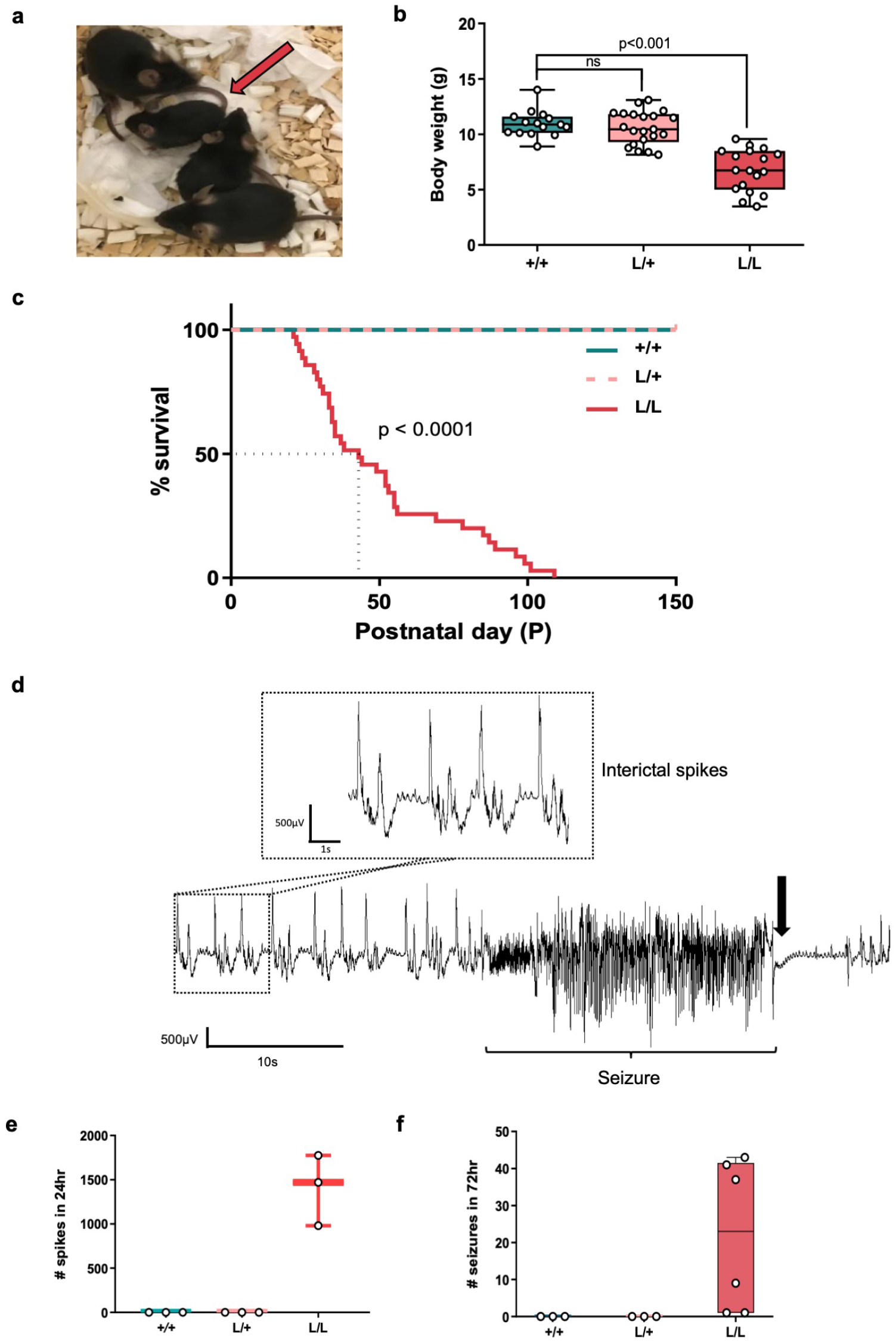
Phenotype of the *Kcnt1* L/L mouse model. **a.** Difference in size at P21 of L/L mice (red arrow) compared to their L/+ and +/+ littermates. **b.** Body weight is significantly reduced in L/L mice compared to their +/+ littermates at weaning age (Kruskal-Wallis test, followed by Dunn’s post hoc analysis; +/+ n=15, L/+ n=21, L/L n=18), data is presented in a box and whiskers plot with maximal and minimal data points. **c.** Life span is shortened in L/L mice with a median survival of 43 days (Kaplan-Meier curve, Log-rank test p-value <0.0001; +/+ n=15, L/+ n=15 and L/L n=35). **d.** Representative ECoG trace of ictal activity and interictal spikes in the L/L mice. Seizures can be preceded by an increase in frequency of acute interictal spikes (square). Spontaneous tonic-clonic seizures correlated with fast, high amplitude signal, followed by electric suppression (black arrow). **e.** Acute high amplitude (>500 μV) spikes are present in the L/L mice, with a median of 1470 spikes in 24 hours (p=0.03, Kruskal-Wallis test, followed by Dunn’s post hoc analysis, n=3 for each genotype) **f.** Seizure frequency over 72 hrs (+/+ n=3, L/+ n=3, L/L n=6).

**Figure 2.**
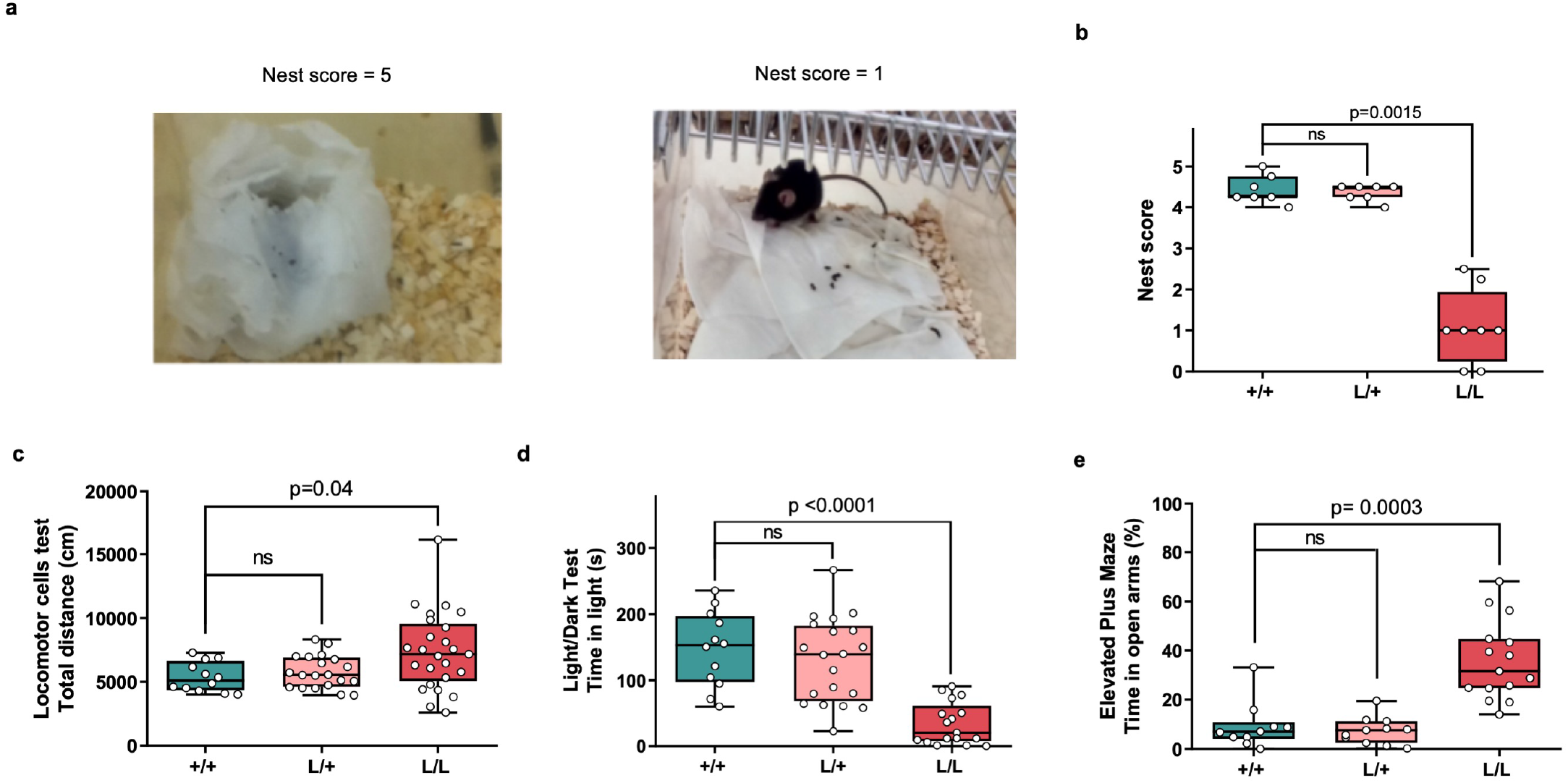
Behavioral profile of the *Kcnt1* L/L mouse model. **a.** Representative images of minimal and maximal scores of nesting behavior. The left image corresponds to a score of 5 (+/+ mouse), while the right image exemplifies a score of 1 (L/L mouse) **b.** Nesting behavior is impaired in L/L mice (+/+ n=7, L/+ n=7, L/L n=8, Kruskal Wallis test with Dunn’s post hoc analysis). **c.** Total ambulatory distance explored in the locomotor cells test. L/L mice are more active compared to L/+ and +/+ (+/+ n=12, L/+ n=20, L/L n=25, Kruskal-Wallis test with Dunn’s post hoc analysis). **d.** L/L mice spend less time in the light compartment during the light/dark box test (+/+ n=12, L/+ n=20, L/L n=17, Kruskal-Wallis test with Dunn’s post hoc analysis) **e.** L/L mice display a preference for the open arms of the elevated plus maze (+/+ n=10, L/+ n=11, L/L n=15, Kruskal-Wallis test with Dunn’s post hoc analysis).

Behavioral abnormalities were also found in the L/L mice. Nest building ability, an identified measure of the general wellbeing in mice (45, 46), has shown to be sensitive to brain lesions, the application of pharmacological agents, and the effects of genetic mutations (47). In mouse models, instinctual nest-building performance is used to simultaneously assess general social behavior, cognitive, and motor performance (48). At P40, the nesting behavior in the L/L mice was markedly impaired compared to their +/+ and L/+ littermates (**Figure 2 a-b**).

Gross motor malfunction and exploratory behaviors were tested on the locomotor cell test and revealed a tendency towards hyperactivity, especially within the first 20 minutes of testing (**Figure 2 c**). The light/dark box test showed L/L mice are prone to anxiety-like traits based on the limited time spent on the light chamber (**Figure 2 d**). The elevated plus maze (EPM) was used to further explore anxiety-like traits. Interestingly, the L/L mice showed a marked preference for the open arms compared to their littermates (**Figure 2 e**), suggesting a reduced fear for open and elevated spaces. Lastly, alterations in social behavior and cognition were tested in L/L mice using the three-chamber social interaction test and Y maze test, respectively, but showed no significant difference compared to +/+ mice (**Extended data figure 1 f-g**).

### ASO-mediated *Kcnt1* reduction rescues the seizure phenotype in L/L mice and improves overall behavior and survival

To determine if the levels of *Kcnt1* mRNA could be reduced in a dose-dependent fashion we tested a mouse-specific *Kcnt1* ASO in adult +/+ mice. At P40, *Kcnt1* ASO was delivered by i.c.v. injection with a range of doses (10, 30, 100, 300 and, 500 μg) and vehicle control-treated animals received an injection of 10 μl of sterile Ca^2+^ and Mg^2+^ free PBS. Two weeks after treatment, the level of *Kcnt1* mRNA was determined using quantitative reverse transcription PCR (RT-qPCR) and compared to that of vehicle control-treated mice. The administration of *Kcnt1* ASO produced a dose-dependent knockdown of *Kcnt1* mRNA in the cortex and the spinal cord (**Figure 3 a-b**). We then determined the specificity of the *Kcnt1*-targetted knockdown by evaluating the expression of the highly-homologous gene Kcnt2 in +/+ mice treated with a bolus i.c.v. injection of 250 μg *Kcnt1* ASO and control ASO (non-hybridizing sequence). *Kcnt1* ASO produced a robust knockdown of *Kcnt1* mRNA in the cortex compared to untreated mice (**Figure 3 c**) but did not affect the expression of *Kcnt2* mRNA (**Figure 3 d**). No significant differences in *Kcnt1* gene expression were found between control ASO treated and untreated mice, confirming that the knockdown observed in *Kcnt1* ASO treated mice resulted from on-target hybridization. Using a pan-ASO antibody we confirmed broad ASO distribution in the mouse brain after a single bolus injection of 75 μg of *Kcnt1* ASO (**Figures 3e**).

**Figure 3.**
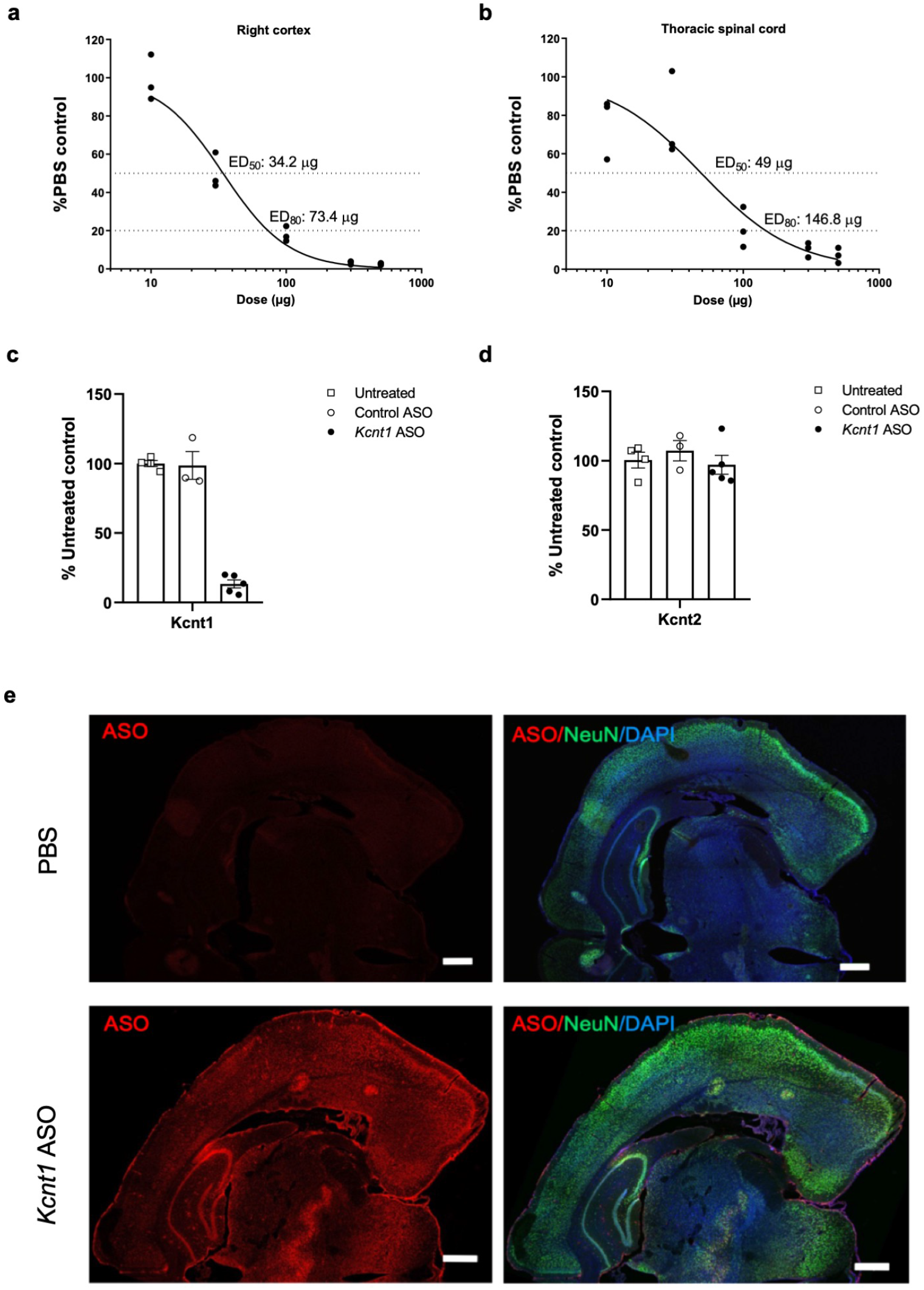
*Kcnt1* ASO produces a dose dependent knockdown of *Kcnt1* mRNA in the mouse CNS. Dose response curves for *Kcnt1* ASO in the brain cortex (**a**) and thoracic spinal cord (**b**) for +/+ mice injected at P40. The tissue was collected 2 weeks after injection and processed for mRNA quantification (n= 3 for each dose, curves were fitted with the Motulsky regression). **c.** The i.c.v. administration of *Kcnt1* ASO reduced the levels of *Kcnt1* mRNA (Kruskal-Wallis test p=0.0027), without affecting the paralogous gene Kcnt2 (Kruskal-Wallis test p=0.4794) (**d**). Mouse cortex was collected 2 weeks after i.c.v. injection for mRNA quantification. Data is presented as bar plots with mean and SEM (untreated n=4, *Kcnt1* ASO n=5, Control ASO n=3). **e.** Distribution of the *Kcnt1* ASO in the mouse brain. Coronal brain sections of +/+ mice treated with *Kcnt1* ASO 75 μg. Tissue was collected 2 weeks after i.c.v. injection and stained with an ASO antibody (red), neuronal marker (NeuN; green) and counterstained with nuclear stain DAPI (blue). *Kcnt1* ASO was found throughout the meninges, hippocampus and cortical layers. Scale bars represent 500 μm.

### ASO mediated *Kcnt1* mRNA reduction rescues the seizure phenotype in L/L mice and improves overall behavior and survival

Survival, seizure frequency, and behavioral markers (nesting, exploratory behavior, anxiety like [light/dark box test, and EPM]) were examined to determine the efficacy of *Kcnt1* knockdown. Adult L/L mice were treated with a single bolus of *Kcnt1* ASO at either ED50 (35μg), ED80 (75μg), or 500 μg. The control group received a dose of 500 μg of control ASO. Treating adult symptomatic L/L mice with a single dose of *Kcnt1* ASO resulted in a significant increase in survival. This effect was dose-dependent and statistically significant across all the tested doses (**Figure 4 b**). In contrast, the animals treated with control ASO displayed a survival consistent with that of untreated L/L mice. Mortality in all treated animals was often related to seizure and presented as status epilepticus.

**Figure 4.**
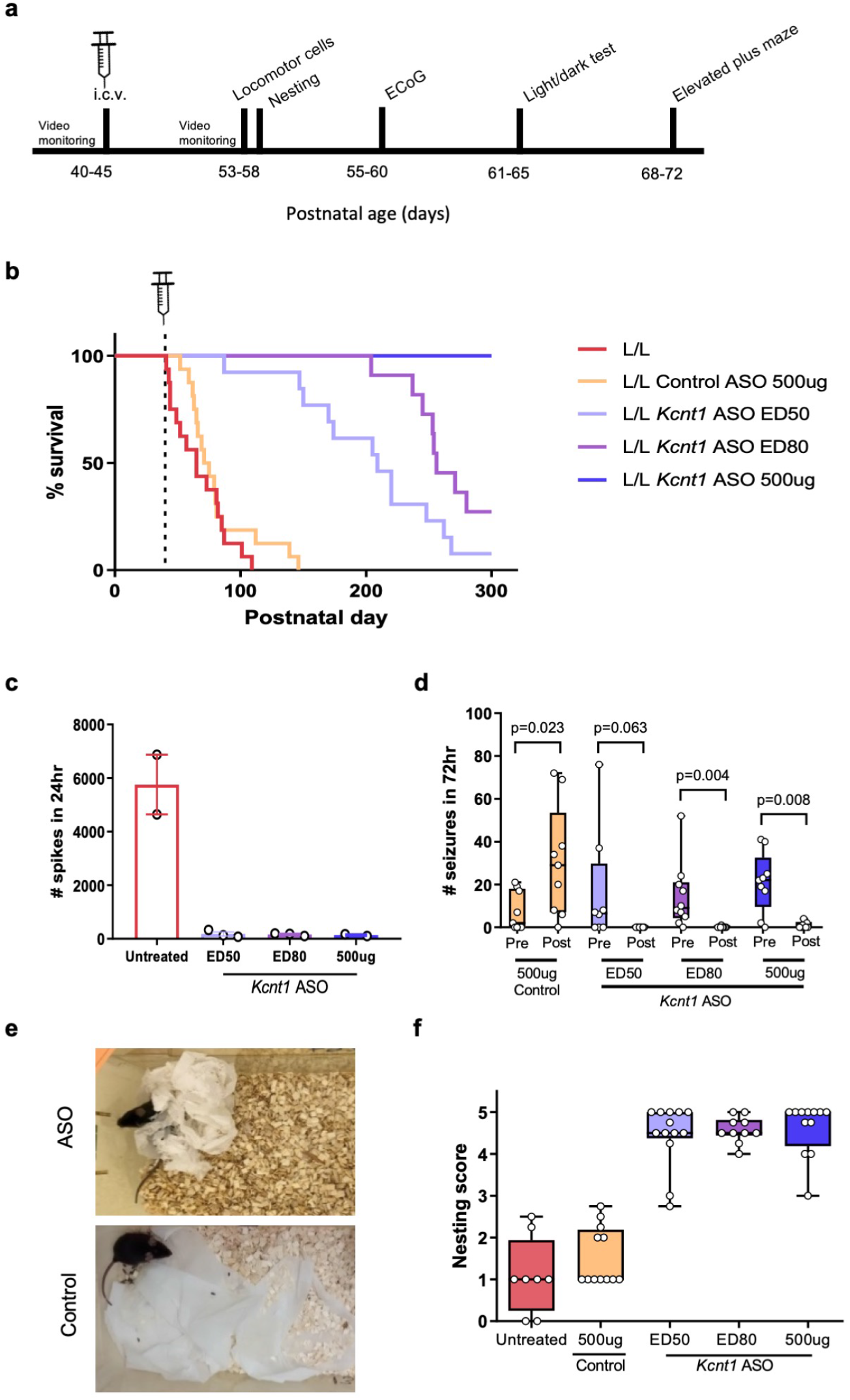
ASO mediated knockdown of *Kcnt1* markedly improves the disease phenotype of adult L/L mice. **a.** Experimental timeline for behavioral studies. **b.** Kaplan-Meier curves shows a dose dependent improvement in survival of adult L/L mice treated with *Kcnt1* ASO (p<0.0001 for *Kcnt1* ASO ED50, ED80 and 500 μg, log-rank test), while mice treated with control ASO showed a survival similar to that of untreated animals (p=0.237, log-rank test, untreated n=16, control ASO n=16, *Kcnt1* ASO ED50 n=13, ED80 n=11, 500 μg n=11). **c.** Acute spike frequency over 24 hrs (n=2 per group). **d.** Seizure frequency was significantly reduced after treatment with *Kcnt1* ASO ED80 and 500 μg. Although ED50 did not reach statistical significance, a trend towards reduction was observed. Treatment with control ASO did not reduce the occurrence of seizures (Control ASO n=9; ED50 n=8; ED80 n=10, 500 μg n=9; seizure frequency was compared using the non-parametric Wilcoxon matched-pairs signed rank test, with Pratt’s method for identical rows). **e.** Representative images of nesting behavior of *Kcnt1* ASO ED80 (top) and Control ASO treated (bottom) L/L mice. **f.** Nesting score of animals treated with *Kcnt1* ASO showed a significant improvement compared to control ASO treated animals (ED50 vs control p=0.0002; ED80 vs control p=0.0015; 500 μg vs control p<0.0001; untreated vs control p>0.9. Kruskal-Wallis test with Dunn’s post hoc analysis. Untreated n= 8, control ASO n=12; ED50 n=13; ED80 n=10, 500 μg n=12).

L/L mice treated with a dose of ED80 and 500 μg of *Kcnt1* ASO had a significant reduction in the total seizure frequency; while the difference in the ED50 group was not significant, a trend towards reduction was observed. In the group that received the control ASO, a significant increase in seizure frequency was seen (**Figure 4 d**). Further, the presence of acute spikes was also reduced in *Kcnt1* ASO treated animals three weeks after treatment (**Figure 4 c**). Age-matched untreated L/L mice were used as controls for ECoG, as the severity of the *Kcnt1* phenotype prohibited meaningful electrical recordings to be made in mice treated with control ASOs. Although the ECoG was not statistical significant, due to the small sample size (n=2-3 for each treatment arm), the trend observed points towards a reduction in the pathological inter ictal signal, which is consistent with the reduced ictal activity observed. During the recording, none of the ASO treated mice presented seizures, while there were 5 seizures recorded in the untreated group. Altogether, these data indicate that the selective knockdown of *Kcnt1* mRNA has an anticonvulsant effect in our model of *KCNT1*-DEE. In addition, mice treated with the *Kcnt1* ASO became less likely to exhibit seizures during routine handling, with no mice presenting seizures during the period of behavioral testing. In contrast, control ASO-treated mice continued to present seizures during routine handling.

Nesting behavior was evaluated 10-13 days after i.c.v. injection. 48-hour after receiving new nesting material, an improvement in nesting behavior was noticed in all animals treated with *Kcnt1* ASO, independent of the dose received (**Figure 4 e-f**). Control treated mice displayed poor nesting behavior, similar to what we observed in untreated L/L mice.

L/L mice treated with *Kcnt1* ASO ED50 and ED80 showed an exploratory behavior similar to that of +/+, while mice that received 500 μg showed a markedly reduced exploratory behavior (**Figure 5 a**). In the light/dark box test, we found no difference in the time spent in the light compartment between mice treated with *Kcnt1* ASO and control ASO, showing an increase in this parameter for all mice that received an i.c.v. injection (**Figure 5 b**). The EPM test showed that independent of the dose received, all *Kcnt1* ASO treated mice spent less time in the open arms of the maze compared to control ASO treated mice (**Figure 5 c**). These data indicate that the ASO silencing of *Kcnt1* not only reduces seizure burden by also normalizes behavioral markers of disease.

**Figure 5.**
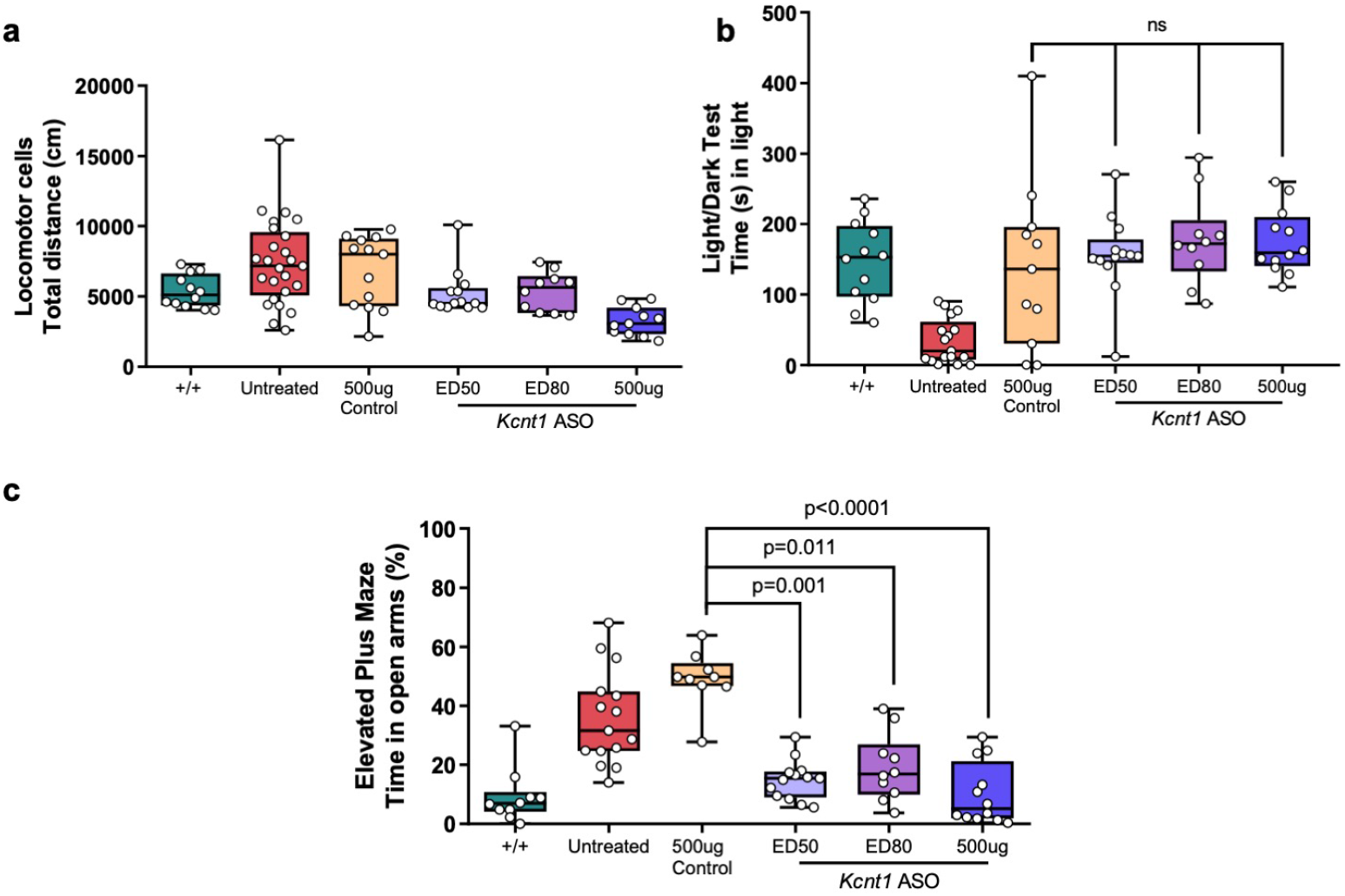
ASO mediated knockdown of *Kcnt1* improves the behavioral phenotype of adult L/L mice. **a.** Total ambulatory distance was reduced in mice treated with *Kcnt1* ASO 500 μg compared to control treated mice (p= 0.0002), but not for mice treated with ED50 or ED80 (p=0.775 and 0.839, respectively) (Kruskal-Wallis test, followed by Dunn’s multiple comparisons, +/+ n=12, untreated n=25, control ASO n=13; ED50 n=13; ED80 n=10, 500 μg n=11). **b.** Time spent in the light compartment during the light/dark box test (Kruskal-Wallis test, followed by Dunn’s multiple comparisons p=0.588; +/+ n=12, L/L n=17, control ASO n=11, ED50 n=13, ED80 n=10, 500 μg n=12). **c.** Time spent in the open arms of the elevated plus maze [Kruskal-Wallis test, followed by Dunn’s post hoc analysis; +/+ n=10; untreated n=15; control ASO n=9; ED50 n=13; ED80 n=10, 500 μg n=12].

### ASO-mediated *Kcnt1* reduction improves general health and allows mating behavior, pregnancy and parental behavior of L/L mice

Due to the severity of the L/L mice phenotype, early mortality and behavioral abnormalities impaired normal mating behavior. To test if mating behavior could be restored, L/L mice received a single dose of 250 μg of *Kcnt1* ASO at P40. Two weeks after injection, male and female mice were set up as breeding pairs or trios. ASO treated dams gave birth to litters of 5 to 8 homozygous pups. Although the first litter was often neglected, both male and female mice displayed effective parental behavior after the second pregnancy. The resulting L/L offspring displayed a phenotype as severe as the homozygous mice obtained from heterozygous breeding if left untreated.

### Neonatal administration of *Kcnt1* ASO is efficacious and well tolerated

To determine if the levels of *Kcnt1* mRNA could be reduced in a dose-dependent fashion at the neonatal period, we tested the *Kcnt1* ASO in newborn +/+ mice. At P2, the *Kcnt1* ASO was delivered by ICV injection with a range of doses (0.5, 1, 1.5, 3, 6 and 30 μg) and vehicle control-treated animals received an injection of 2 μl of sterile Ca^2+^ and Mg^2+^ free PBS. Two weeks after treatment, the level of *Kcnt1* mRNA was determined and compared to that of vehicle control injected mice. The early administration of *Kcnt1* ASO produced a dose-dependent knockdown of *Kcnt1* mRNA in the mouse cortex (**Figure 6 a**).

**Figure 6.**
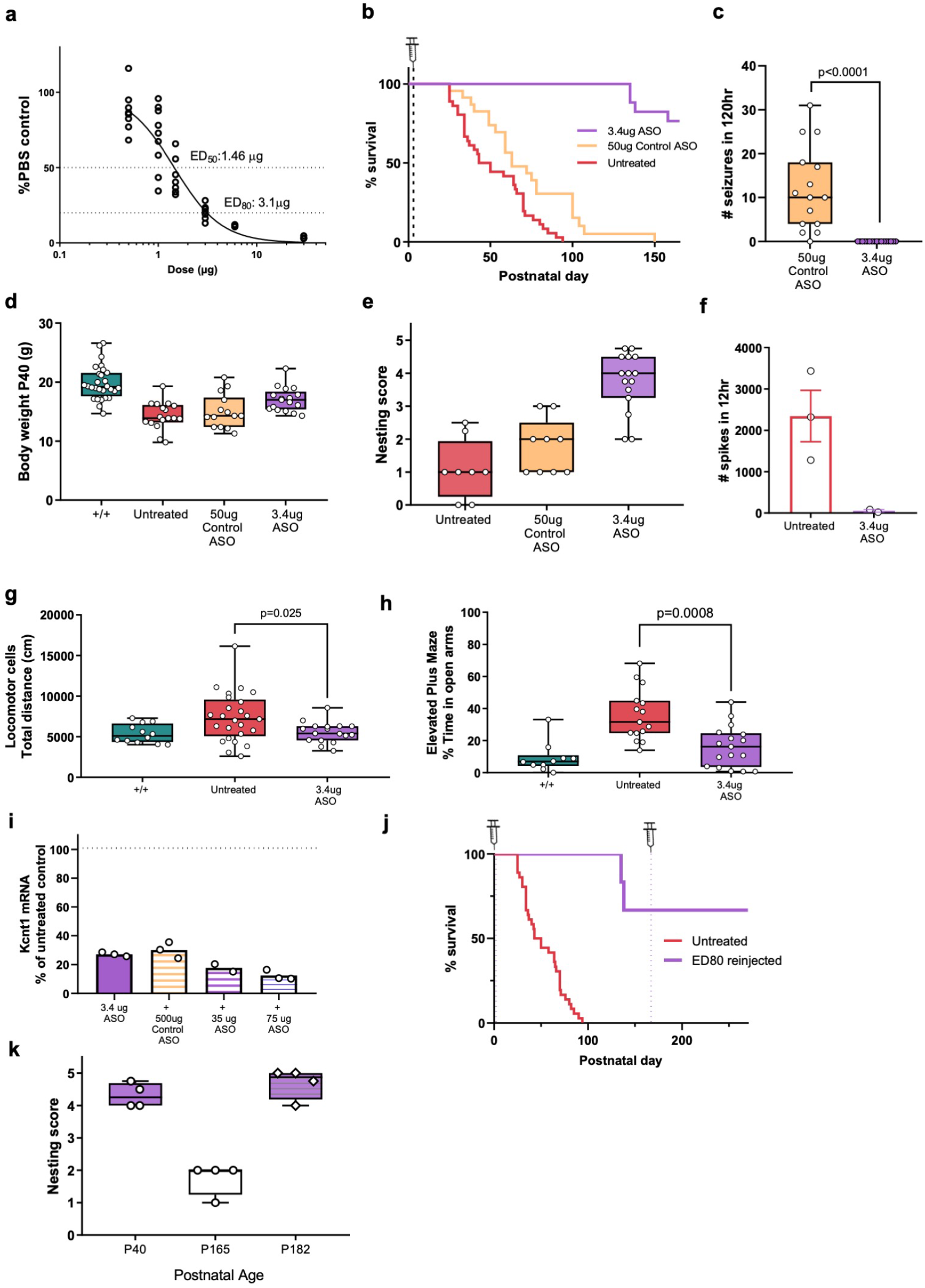
Neonatal administration of *Kcnt1* ASO is safe and effective in the L/L mouse model. **a.** Dose response curve for +/+ mice injected at P2. Brain cortical tissue was collected 2 weeks after injection (n=3-11 per dose, the curve was fitted with the Motulsky regression). **b.** Kaplan-Meier curve for neonatal (P2-P3) administration of *Kcnt1* ASO in L/L mice. Lifespan of *Kcnt1* ASO treated mice is prolonged compared to Control ASO treated mice (p<0.0001). 50 μg of control ASO produced a smaller but significant improvement in survival (p=0.003) (untreated n= 36; control ASO n=23; *Kcnt1* ASO 3.4 μg n= 17). **c**. Seizure frequency was reduced after neonatal administration of *Kcnt1* ASO when compared to control treated (50 μg) mice (p<0.0001, Mann Whitney test, control ASO 50 μg n=15, *Kcnt1* ASO 3.4 μg n=17). **d.** Weight at P40 of neonatal treated L/L mice improved compared to control treated mice (p=0.016, Mann-Whitney test, +/+ n=28, untreated n=17, control ASO 50 μg n=14, *Kcnt1* ASO 3.4 μg n=17). **e.** Nesting behavior improved in L/L mice treated with *Kcnt1* ASO (p =0.0001, Mann Whitney test, untreated n=8; control ASO 50 μg n=9; *Kcnt1* ASO 3.4 μg n=15). **f.** Spike frequency in neonatal L/L mice treated with *Kcnt1* ASO. **g.** Total ambulatory distance was reduced in mice treated with *Kcnt1* ASO 3.4 μg compared to untreated mice (Mann Whitney test, +/+ n=12, untreated n=25, *Kcnt1* ASO 3.4 μg n=17). **h.** Time spent in the open arms of the elevated plus maze (Mann Whitney test; +/+ n=10, untreated n=15, *Kcnt1* ASO 3.4 μg n=17). **i.** A second injection of *Kcnt1* ASO at P30 was tolerated in +/+ mice with a further reduction of *Kcnt1* mRNA compared to a single injection at P2. Cortical tissue was collected at P42 *[Kcnt1* ASO 3.4 μg n =3; *Kcnt1* ASO 3.4 μg + Control ASO 500ug n= 3; *Kcnt1* ASO 3.4 μg + *Kcnt1* ASO 35 μg n=2; *Kcnt1* ASO 3.4 μg + *Kcnt1* ASO 75 μg n=3). **j.** Survival curve for L/L mice treated with a second dose of ED80 ASO at P167 (dotted line) (Untreated n= 36; ED80 reinjected n=4). **k.** Nesting score of L/L mice reinjected with ED80 ASO. After a 3.4 injection at P2, nesting behavior was present at P40 but had declined by P165. A second injection of 75 ug ASO rescued the nesting behavior (n= 4).

Then we examined whether the early administration of *Kcnt1* ASO could prevent the development of a disease phenotype in the L/L model. At P2-P3, L/L mice received a single i.c.v. bolus injection of 3.4 μg (ED80), while control ASO mice received an injection of 50 μg. *Kcnt1* ASO treated mice showed extended survival, with the first deaths being observed at P135 (**Figure 6 b**) and displaying approximately 25% mortality by P150. Similarly, control ASO treated mice also showed a shorter yet significant improvement in survival (median survival of 63 days compared to 46.5 for untreated mice). Seizure frequency improved significantly in mice treated with *Kcnt1* ASO and interictal ECoG activity was also reduced (**Figure 6 c and f**). Bodyweight was measured at P40 and was increased for *Kcnt1* ASO L/L mice compared to that of controls but overall, body weight was still lower compared to +/+ mice (**Figure 6 d**). Similar to adult treated L/L mice, behavioral markers including nesting and hyperactive exploratory activity (locomotor cells) and EPM showed improvement compared to untreated mice (**Figure 6 e, g, h**).

Considering the likelihood that an ASO therapy for *KCNT1*-DEE in patients would require chronic ASO administration, we tested if a second dose was tolerated. A small group of +/+ mice received a 3.4 μg dose of *Kcnt1* ASO at P2 and a second injection of either *Kcnt1* ASO (35 or 75 μg) or control ASO (500 μg) at P30. The procedure was tolerated in all treatment groups. By P42, the administration of a second dose of 75 μg *Kcnt1* ASO showed a further reduction of *Kcnt1* mRNA compared to mice receiving control ASO on the second injection (**Figure 6 i**).

In a small subset of L/L mice (n=4) treated at P2, we tested if a second dose of *Kcnt1* ASO (75 μg) could be effective after the animals have become symptomatic (i.e. nesting behavior was deficient and seizures at handling were observed in 1 mouse). In these cases, a further extension in survival was observed (**Figure 6 j**) and nesting behavior was rescued (**Figure 6 k**).

### ASO-mediated *Kcnt1* reduction in +/+ mice is well tolerated

To evaluate the consequence of *Kcnt1* reduction, we treated adult +/+ mice with 500 μg *Kcnt1* ASO and performed an array of behavior tests. ASO-mediated knockdown of *Kcnt1* mRNA in +/+ mice did not affect their nesting behavior (**Extended data Figure 2 a**) and was consistent with a previously reported study on a *Kcnt1* knockout (KO) mouse model (104). The exploratory behavior in the locomotor cell test was similar to that of control-ASO treated mice and although the total ambulatory distance explored displayed a tendency towards reduced activity, this difference did not reach statistical significance (**Extended data Figure 2 b**). Similarly, a tendency to spend less time in the light chamber was observed (**Extended data Figure 2 c**).

On the EPM, +/+ *Kcnt1* ASO treated mice showed a behavior similar to that observed in L/L *Kcnt1* ASO treated mice, with a reduction in the time spent in the open arms **(Extended data Figure 2 d).** In the three-chamber social interaction test, no difference was found between control and *Kcnt1* ASO treated mice in the time spent with an intruder mouse **(Extended data Figure 2 e).** Finally, to further explore changes in memory, the mice were tested in the Y maze. A reduction in the median time spent in the novel arm was observed in *Kcnt1* ASO treated mice compared to control **(Extended data Figure 2 f).**

## DISCUSSION

In this study we investigated the potential of an ASO-based gene silencing approach as a potential therapy for *KCNT1*-DEE. We have shown a gene-specific and dose-dependent knockdown of *Kcnt1* mRNA achieved with the *Kcnt1* ASO in L/L mice which significantly improved survival, epilepsy, behavioral comorbidities and complete rescue of mating.

We first developed a rodent model and established disease biomarkers that could be used for therapeutic screening. While heterozygosity for a gain of function variant in *KCNT1* is sufficient to result in a disease state in patients with gain of function *KCNT1* variants, this was not replicated in our mouse model. L/+ mice did not display an epileptic phenotype, increased susceptibility to seizures or pathologic behavioral changes. This is not completely unexpected for mouse models of genetic epilepsy demonstrate considerable phenotypic variability based on factors including strain (49) and sub-strain (50) used. In contrast, homozygous mice presented a robust epilepsy phenotype, behavioral deficits and a progressive deteriorating course resulting in reduced survival, thus presenting a valuable tool for drug screening.

Within 2 weeks of bolus administration, ED80 and 500 μg doses of *Kcnt1* ASO produced a significant reversal of the epileptic phenotype. While dosing at ED50 showed only a trend towards seizure reduction, which could have been influenced by the sample size on this treatment group. A clear dose-dependent effect was observed for survival of adult L/L treated mice. Overall, the effectiveness of the ASO approach in controlling seizure activity confirms that brain hyperexcitability in the disease model is driven by *Kcnt1* gain of function and therefore by reducing the excessive channel activity the seizure threshold can be modified.

A critical concern shared among neurodevelopmental conditions and neurodegenerative disorders is whether disease progression can be stopped and the cognitive capacity preserved or improved once the pathological process has begun. Importantly, we have shown here that the downregulation of *Kcnt1* can be safe and effective in both neonatal (presymptomatic) and adult (symptomatic) mice by not only in controlling spontaneous seizures but also providing disease modifying effects as indicated by improvements in cognition and behavior.

We also asked if the down-regulatory ASOs had any harmful effects derived from excessive knockdown. Studies on *Kcnt1* KO mice have reported some mild behavioral and cognitive alterations (51, 52). Here we show that the administration of a high dose of *Kcnt1* ASO to adult (500 μg) and neonatal (30 μg) +/+ mice did not result in serious adverse events. Treated adult mice showed a nesting behavior similar to that of untreated mice, and consistent with the previously reported phenotype of *Kcnt1* KO (52).

A reduced exploratory behavior in the open field was reported in the *Kcnt1* KO model characterization published by Bausch *et al* (51). Interestingly, while the initial exploratory activity of wild-type ASO treated mice was similar to that of the control treated group, we did identify a trend that indicated an overall reduced exploratory activity, which further supports the idea that Kcnt1 is important for the cognitive process of exploring and adapting to new environments (51). More importantly, a completely contrasting behavior (hyperactivity) in the L/L disease model further supports this observation (51).

RNA-targeted ASO technology is an innovative therapeutic modality that has already achieved clinical success as an effective treatment for rare neurogenetic conditions like spinal muscular atrophy (43), Duchene’s muscular dystrophy (44) and hereditary transthyretin amyloidosis (53). The FDA approval of nusinersen marked an important milestone for ASO technology.

Nusinersen has demonstrated the disease-modifying properties and fulfilled the promise of precision medicine, opening a path for the further development of other RNA based therapies for neurogenetic diseases (38). Although CNS targeting ASOs require direct intrathecal delivery to be effective, this invasive approach is compensated by the extended half-life in the target tissue, and the wide brain distribution and high cellular uptake (54–57) which allows for less frequent dosing. In addition, direct delivery to the CSF compartment precludes the ASO from being distributed to the rest of the body, limiting the occurrence of adverse effects (54).

Genetic neurodevelopmental conditions, including DEE are devastating conditions that lack effective therapies. The unravelling of the genetic architecture of DEE has provided an exciting opportunity to develop precision medicines. Therapeutics that use RNA targeting molecules are well positioned to directly address the underlying cause of DEE including seizures and comorbid pathologies. In this study, we demonstrated that ASO-mediated reduction of *Kcnt1* is safe and has a disease-modifying effect in a mouse model of *KCNT1*-DEE. These findings provide crucial evidence to support the development of ASO based therapies for refractory epilepsies and developmental disorders that can contribute to reducing the impact of neurogenetic disorders.

## METHODS

### *Kcnt1* knock-in mouse model

Mice were housed in temperature-controlled rooms (≈22°C) on a 12 h dark/light cycle, weaned at ≈ postnatal day 21 (P21), and maintained with rodent diet and water available *ad libitum*. Male and female mice were used for all experiments. Mice were housed in individually ventilated cages (IVC, Tecniplast, Sealsafe plus mouse IVC green line) in groups of up to five animals. Mice that underwent ECoG electrode implantation, seizure monitoring, and nesting behavior assessment were housed individually during recovery of surgical procedures and for the duration of the test.

### Mouse model generation

The L/L knock-in mouse line was generated at the University of Adelaide using CRISPR/Cas9 to insert a single nucleotide mutation at position c.2714, changing C>T. The *Kcnt1* p.P905L mutation is homologous to the *KCNT1* p.P924L mutation. *Kcnt1* p.P905L founders were generated as described previously (58). C57BL/6J zygotes were injected with 50 ng/μl of sgRNA (TGCATGAACCGCATGTTGGA), 100 ng/μl of Cas9 mRNA and 100 ng/μl of a singlestranded oligonucleotide repair template (Ultramer DNA from Integrated DNA Technologies). The oligonucleotide repair template sequence was:

GTTTGGAAAGAGCCAGAGAGTAGCTGTCCTTGGCACGGAACTGCATGAACCGCATG <u>TTCGAA</u>aGGTGTGTGAGCTCCGTGGTGATGCTGAGACTGGGGAAAAGCCTGAGGGA GGATGATCG.

Injected embryos were transferred to pseudopregnant females for further development. Pups were genotyped by PCR and the intended C>T mutation was confirmed by Sanger sequencing. The colony was maintained on the C57BL/6J background and initially backcrossed to wild-type (+/+) mice to expand the colony with subsequent heterozygous intercross carried out to obtain wild-type (+/+), heterozygous (L/+) and homozygous (L/L) mice. Where possible *+/+* littermates were used as controls. When unavailable or as “intruders” for behavioral paradigms (specifically the three chamber social interaction test), C57BL/6J wild-type mice were purchased from the Animal Resources Centre (Canning Vale, WA, Australia).

For experiments with neonatal mice: C57BL/6J wild-type pregnant mice were purchased from the Animal Resources Center (Canning Vale, WA, Australia). L/L litters were obtained from the crossing of *Kcnt1* ASO-treated L/L breeders.

### Identification and genotyping

Animals were toe-clipped at postnatal days 8 to 12 (P8 - P12) for identification and the tip of the tail was biopsied for genotyping. DNA extraction was performed using the REDExtract-N-Amp Tissue PCR Kit (Sigma, St Louis, MO, USA).

The following PCR primers were used to amplify exon 24 of *Kcnt1*:

Forward: 5’-CCACCCAGTTATGACCACAG-3’
Reverse: 5’-GCTGTAGGTATCTGTTAGCAG-3’

PCR products were digested with the restriction enzyme BstBI (New England BiolLabs Inc, Ipswich, MA, USA) and the products were separated through electrophoresis on a 2% agarose gel stained with GelRed (Biotium, Fremont, CA, USA). The wild-type allele generated a single fragment of 460 bp and the mutant allele generated two fragments of 276 and 184 bp.

### Mouse model phenotypic validation

#### Spontaneous behavioral seizures video recording

Mice older than P21 were housed individually in a 19.56 x 30.91 x 13.34 cm Thoren # 9 Small mouse II cage (Thoren Caging System, Hazleton, PA, USA) and monitored continuously for up to 5 days. Video recording was done using the Vivotek video server (VS8102) connected to an infrared day and night digital color camera (EVO2; Pacific Communications). Following recording, the videos were played back on a computer at 8-13x speed using the VLC media player (Paris, France). Seizures that presented with any of the following behaviors were counted as an event: a) clonic seizure in a sitting position; b) clonic and/or tonic–clonic seizures while lying on the belly c) pure tonic seizures; d) clonic and/or tonic–clonic seizures while lying on the side and/or wild jumping. A tonic clonic seizure lasting more than 3 minutes was considered to be prolonged, and a seizure with a duration of more than 5 minutes was considered to be a status epilepticus.

#### Susceptibility to induced seizures

##### Thermogenic seizure assay

Mice between the ages of P18-P21 were placed in a container heated to constant 42°C for a maximum of 20 minutes. The latency to a first tonic-clonic seizure was recorded.

##### Pentylenetetrazol induced seizures

Mice between the ages of P30-P40 were injected subcutaneously (s.c. injection) with the GABA antagonist pentylenetetrazol (PTZ; 80mg.kg^-1^; Sigma, St. Louis, MO, USA) dissolved in 0.9% sterile saline solution (Pfizer, NY, USA). The animals were then monitored for a maximum of 45 minutes and latencies to a minimal (first tonic-clonic) and maximal (tonic hindlimb extension) seizures were recorded.

##### Loxapine induced seizures

Mice between the ages of P30-P40 received a peritoneal injection (i.p. injection) with the antipsychotic drug Loxapine (LOX; 100mg.kg^-1^; Sigma, St. Louis, MO, USA) dissolved in 0.9% sterile saline solution (Pfizer, NY, USA) and 25% Dimethyl Sulfoxide (DMSO, Sigma, St. Louis, MO, USA). The animals were then monitored for a maximum of 1 hour and the latency to a first tonic-clonic seizure was recorded.

In the three assays, animals were euthanized at the end of the experiment.

#### Behavioral profile

Behavioral tests were conducted between 9:00 A.M. and 6:00 P.M., under similar lighting conditions for each task. The mice were transferred to the behavior testing room at least 30 min before the test. All mice were subjected to each behavioral test with a 5-7-day interval between each task.

##### Nesting behavior

Two facial tissues (18 x 18,5 cm;1.75-1.9 g) (Austwide Paper products, VIC, Australia) were provided as nesting material two hours prior to the onset of the dark phase (5-6pm). The quality of the nest was assessed 48h after. A score from 0 to 5 was given by adapting the scoring system of Hess et al (45).

##### Locomotor cells

Mice were placed in the center of a 27.31 x 27.31 x 20.32 cm covered chamber (Med Associates Cat# ENV-510S-A) for 30 min and their activity was monitored by a 48 channel infrared (IR) controller in 5 min bins and analyzed with the Activity Monitor Software Version 7 (Med Associates, Fairfax, VT, USA).

##### Light/Dark Box

The light–dark apparatus consisted of a 27.31 x 27.31 x 20.32 cm chamber (Med Associates Cat# ENV-510S-A) divided into dark and light compartments of equal size by the insertion of a black Perspex box. The box contained a small opening in the middle allowing the mouse to move between the compartments. The light chamber was brightly illuminated to 750 Lux by LED light lamps. Each mouse was placed in the center of the dark compartment and allowed to freely explore the box for 10□minutes. The latency to move into the light compartment as well as the amount of time spent in each side were automatically recorded and analyzed with the Activity Monitor software version 7 (Med Associates, Fairfax, VT, USA).

##### Elevated plus maze (EPM)

The elevated plus-maze apparatus was made of light-colored Perspex and consisted of two open arms (5 x 30 cm) and two enclosed arms (5 x 30 x 14 cm) extending from a central area (5 × 5 cm). A raised lip (2.5 mm high, 5 mm wide) around the open arms was placed to reduce the likelihood of mice falling off the maze. The maze was elevated 60 cm above the floor. Each mouse was placed in the central area, facing an open arm. Their activity in the maze was video recorded for 10 min. The time spent on each arm of the maze and in the center area was measured, and arm entries were counted using the TopScan software (CleverSys, Reston, VA, USA).

##### Three chamber social interaction

The test was conducted in a box (43 x 39 x 11 cm) made of transparent plastic and divided into three chambers; the middle section was 8 x 39 cm and the two lateral sections were 16 x 39 cm each. The chambers were connected by rectangular openings in the middle. Metal mesh cages of 16 x 10 x 11 cm were placed in the two side chambers. The test consisted of two phases: habituation and trial phase. During the habituation period, the mouse was placed in the central chamber and allowed to explore freely all chambers for 10 min. After that, a preference for a chamber (left vs right) was assessed and an unfamiliar mouse of similar age and size was placed in the metal cage of the side chamber of least preference. For the trial period, the mouse was again allowed to explore all chambers for 10 min. The activity during both habituation and trial period was video-recorded and the amount of time spent in each chamber and near the cages was analyzed using TopScan software (CleverSys, Reston, VA, USA).

##### Y maze

The Y maze was made of light-colored Perspex and consisted of three 7.5 x 30 x 14 cm arms separated by a 120° angle and containing a visual cue at the end. The test was performed in two phases: a training session and a trial session. During the training session the mouse was placed in the far end of one arm (home arm), facing away from other arms, and was allowed to explore this and one other arm (familiar arm) for 10 min. During this period, the third arm (novel arm) was not accessible. After a 1-hour interval, the mouse was again placed in the home arm but allowed to explore all three arms for 5 min (trial session). The test was video recorded, and the amount of time spent in each arm was analyzed using TopScan software (CleverSys, Reston, VA).

### Surgical procedures

#### ECoG Electrode implantation surgery

Mice were anaesthetized with 3-4 % Isoflurane (IsoFlo; Abbott Laboratories, Abbott Park, IL, USA) for induction and 2% for maintenance. A subcutaneous injection of meloxicam (1 mg/kg) dissolved in 0.9% saline was given prior surgery. Then, the mouse was placed in a stereotaxic apparatus (Kopf Instruments, Tujunga, CA, USA), after the scalp was shaved, sterilized with 80% ethanol and infiltrated with Lignocaine (1% ampules, Pfizer). A 1 cm long incision was made on the scalp and the skull was cleaned with 3% hydrogen peroxide solution (Sanofi). Four burr holes were drilled into the skull and three screws were placed into the holes and used as epidural electrodes (Cat #8403, Pinnacle Technology, Lawrence, KS, USA) (1 reference electrode and 2 recording electrodes). A ball of silver wire was used as ground electrode. All electrodes were connected to a headmount (Cat # 8201, Pinnacle Technology, Lawrence, KS, USA) and affixed to the skull with methyl methacrylate dental cement (Cat # 1255710; Henry Schein Inc, Indianapolis, IN; Lang Jet Denture Repair Acrylic). Then, mice recovered in a warming pad at 30 °C until fully awake.

#### ECoG recording and analysis

Mice were allowed to recover for 3-5 days before recording. The mice were connected while awake to a mouse pre amplifier (Cat # 8406-SE) and an amplifier (Cat #8204 and 8206, Pinnacle Technology, Lawrence, KS, USA). Brain cortical activity was sampled on Sirenia (Pinnacle Technology, Lawrence, KS, USA) at 250 Hz for 25 hours, signal was band-pass filtered at 0.5 to 40Hz. ECoG signal was analyzed post acquisition using ClampFit 10.7 (Molecular Devices), a spike was identified when the amplitude was 2.5 times greater than the baseline activity and duration was shorter than 80 ms. An investigator blinded to the treatment groups/genotype reviewed and counted spike activity.

##### Intracerebroventricular (i.c.v.) injections

###### Adult mice

P40-P45 mice, weighing at least 10g, were anesthetized using Isoflurane (IsoFlo; Abbott Laboratories, Abbott Park, IL, USA) at a concentration of 4-5% mixed in O_2_ (vol/vol) for induction, and 2% for maintenance. A subcutaneous injection of meloxicam (1 mg/kg) dissolved in 0.9% saline was given prior surgery. The mice were positioned in a stereotaxic apparatus (myNeuroLab, Leica, Germany) and the scalp was cleaned with 80% Ethanol. Then, the skin was infiltrated with Lignocaine (1% ampules, Pfizer) and a 1 cm incision was performed. The skull was cleaned with 3% hydrogen peroxide solution to expose the bone sutures and a burr hole was drilled at 0.8 mm lateral and 0.3 mm posterior to Bregma. The tip of a 33G internal infusion cannula (Plastics One Cat #C315I/SPC) was advanced to −3.0 mm from the skull surface to reach the right lateral ventricle. The cannula was connected to a 0.5 ml glass syringe (SDR, Sydney, Australia) and an infusion pump (legato 210/210p syringe pump, Kd Scientific, Holliston, MA, USA). A total volume of 10 μl of ASO or vehicle (sterile Ca^2+^ and Mg^2+^ free PBS) was delivered at a rate of 0.5 μl/s. One minute after completing the infusion, the cannula was withdrawn and the skin was closed with 4-0 polyglactin 910 absorbable suture (coated Vycryl, Ethicon Inc, Cornelia, GA, USA). Following surgery, the animals recovered on a Thermacage (Datesand, Ltd, Manchester, UK) until active and were then returned to their home cage.

###### Neonatal mice

P2-P3 pups were cryo-anesthetized for 3 minutes. The scalp was cleaned with 80% ethanol and free hand injections were performed using a 32 G needle Hamilton syringe (10μl, Hamilton, Reno, NV, USA). Using lambda and the right eye as anatomical references, the needle was inserted midway between these two points and advanced −2.0 mm ventral from the skin surface (technique adapted from Kim *et al* (59)). A total volume of 2 μl of ASO or vehicle was injected into the right lateral ventricle. Pups were gently warmed until skin color returned to pink and rolled over dirty bedding before being returned to their home cage. Maternal behavior was monitored for the following 10 minutes to ensure the pups were attended and returned to the nest.

### ASO synthesis

The *Kcnt1* ASO and control ASO (non-hybridizing sequence) were synthesized and screened *in vitro* by Ionis Pharmaceuticals (Carlsband, CA, USA) as previously described (60, 61). Both molecules are 20bp long and have the following structure: 5 modified nucleotides with 2’-MOE modifications at the 5’ and 3’ end, and a central gap (gapmer) of 10 unmodified oligodeoxynucleotides. A PS backbone was used to enhance nuclease resistance. The specific sequence for each ASO is listed below:

*Kcnt1* ASO: 5’-GCTTCATGCCACTTTCCAGA-3’
Control ASO: 5’-CCTATAGGACTATCCAGGAA-3’

ASOs were solubilized in sterile Ca^2+^ and Mg^2+^ free Dulbecco’s Phosphate Buffered Saline (PBS) (Sigma, St Louis, MO, USA), centrifuged, and filtered through a 0.22 μm filter (Sigma, St Louis, MO, USA). A stock solution with a concentration of 50 mg/ml was stored at −20°C. Then, ASOs were further diluted to the desired concentration in sterile Ca^2+^ and Mg^2+^ free PBS (Sigma, St Louis, MO, USA) immediately before injection.

### Quantification of mRNA levels

Tissue harvest: Two weeks after ASO administration, mice were deeply anesthetized with 4-5% Isoflurane (IsoFlo; Abbott Laboratories, Abbott Park, IL, USA), before being decapitated. After the brain was removed from the skull, the cerebellum, right and left cortex were dissected. The spinal cord was harvested using hydraulic extrusion as previously described (62) and a 1 cm piece from the low thoracic/lumbar cord was extracted. All dissected tissue was snap frozen in liquid nitrogen, and stored at −80°C until RNA preparation.

Quantitative gene expression analysis: Total RNA was isolated from the mouse right cortex and spinal cord using TRIzol reagent (Cat #15596026, Thermofisher, Waltham, MA, USA) according to the manufacturer’s protocols. Contaminating genomic DNA was removed with DNAse treatment (DNA-free Reagents; Ambion/Life Technologies, Carlsband, CA, USA). Primer sequences were designed to identify the following mRNA:

*Kcnt1*
Forward: 5’-TCTTCCCTTTCTCAGGTCCAGG-3’
Reverse: 5’-AGGGAGAAGTTGAACAGCCG-3’
Kcnt2
Forward: 5’-AAGGCTGAGCAGAAAAGGGC-3’
Reverse: 5’-TCATTTCATCGTAGCCCACCG-3’
Rpl32
Forward: 5’-GAGGTGCTGCTGATGTGC-3’
Reverse: 5’-GGCGTTGGGATTGGTGACT-3’

For quantitative reverse transcription and polymerase chain reaction (RT-qPCR), oligo deoxythymidylic acid (oligo-dT) primed cDNA was synthesized from 500 ng of total RNA using Murine Moloney Leukaemia Virus Reverse Transcriptase (Promega, Madison, WI, USA). RT-qPCR was performed on the ViiA 7 Real-Time PCR System (Applied Biosystems/Thermofisher, Foster City, CA, USA) using GoTaq qPCR master mix (Promega, Madison, WI, USA) according to the manufacturer’s protocols. Relative gene expression values were obtained by normalization to the reference gene Rpl32 using the −2ΔΔCt method, where −2ΔΔCt = ΔCt sample–ΔCt calibrator as previously described (63).

### Immunohistochemistry

Tissue preparation: Two weeks after ASO administration, mice were anesthetized with sodium pentobarbitone (80 mg/kg, lethal dose, i.p. injection) (Ilium, Troy Laboratories, Smithfield, Australia) and transcardially perfused with 0.1 M phosphate buffer (PB) followed by a 4% paraformaldehyde solution (PFA; pH7.4). Brains were removed from the skull and stored in 20% sucrose solution overnight at 4°C, and then frozen in isopentane cooled in liquid nitrogen before being sectioned coronally at 20 μm thickness with a cryostat (Leica Microsystems, Wetzlar, Germany). Sections were mounted on Superfrost plus slides (Thermofisher, Waltham, MA, USA) and stored at −20 °C until used for immunostaining.

Immunostaining: Sections were air-dried for 1 hour at room temperature and then blocked for 1 hour in a humidified chamber, with a mixture of 10% normal goat serum, 0.3% Triton X-100 (Sigma, St Louis, MO, USA) in PB. The blocked sections were incubated overnight with primary antibodies: polyclonal rabbit anti-ASO 1:7500 (Ionis Pharmaceuticals); polyclonal guinea-pig anti-NeuN 1:500 (Millipore; Cat #ABN90-clone A60 [MAB377]); After washing in PB, the sections were incubated for 2 h with secondary antibodies: goat-anti-rabbit Alexa-647 1:500 (Thermo-fisher; Cat#A31573) and donkey-anti-guinea pig Alexa 488 1:500 (Thermo-fisher; Cat# A-11073). To identify the nuclei, sections were then stained with DAPI (Sigma-Aldrich). All incubations were conducted at room temperature. Slides were covered with Prolong Gold Anti-fade (Invitrogen) and stored at −30 °C. Confocal image stacks were acquired on a Zeiss LSM 780 microscope equipped with a 20 x/0.8 NA lens. Z-stack images were acquired within Nyquist sampling parameters and de-convolved using Huygens 4.4.0 software (Scientific Software Imaging).

### Statistical analysis

The latencies for the seizure susceptibility tests were plotted as Kaplan Meier survival curves and tested for significance using the Log rank (Mantel Cox) test. In all the survival plots, animals that did not reach the end point by 40 minutes (20 minutes for the thermogenic assay) were marked as censored. Dose-response curves were fitted with the Motulsky regression. For all other measures, Fischer test, Mann–Whitney U test, nonparametric one-way and two-way ANOVA with post hoc analysis were used accordingly. Statistics were computed using GraphPad Prism, San Diego, CA, USA. For all tests, statistical significance was set at a p < 0.05.

### Study approval

All animal experiments were approved by the animal ethics committee of the Florey Institute of Neuroscience and Mental Health (protocols #14-026, 16-061, 16-062 and 17-014) and were performed in accordance with the guidelines of the National Health and Medical Research Council Code of Practice for the Care and Use of Animals for Experimental Purposes in Australia.

## Acknowledgements

We thank Mr Brett Purcell and Mr Travis Featherby for their assistance with behavioral experimental setup and Ms Taryn Knight, Ms Ana Hudson, Mr Daniel Drieberg and Mr Juan Raffin for their assistance with mouse colony maintenance. We also thank Ms Lisa Drew and Dr Svenja Pachernegg for their assistance with behavioral tests and RT-qPCR.

## EXTENDED DATA

**Supplementary figure 1.**
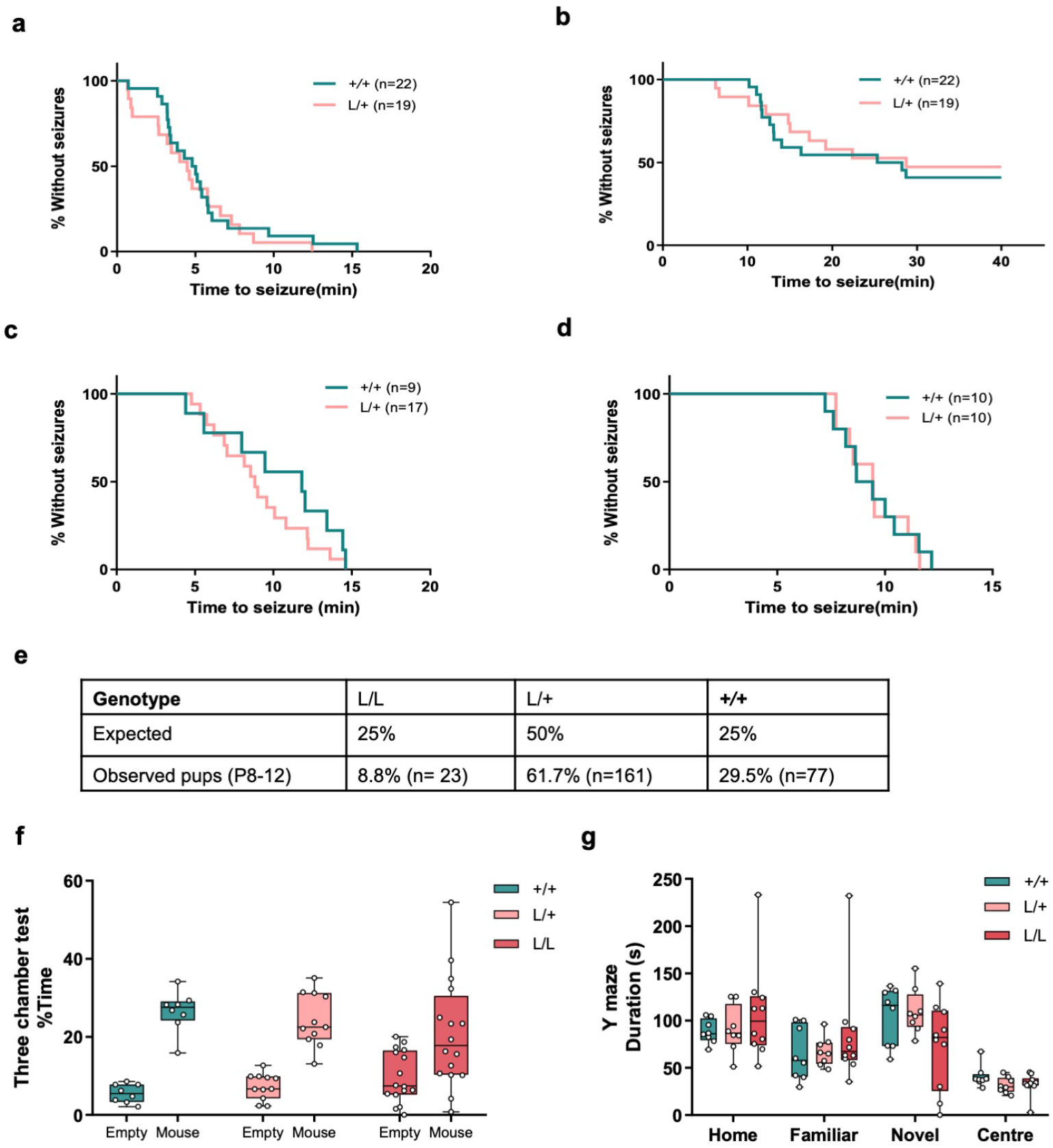
Extended phenotypic characterization of the *Kcnt1* p.P924L mouse model. Chemo-convulsant challenge with PTZ showed no increased susceptibility in L/+ mice for a first tonic-clonic seizure (**a**) (Kaplan-Meier curve, Log-rank test p= 0.55) and a severe seizure with tonic hindlimb extension (**b**) (Kaplan-Meier curve, Log-rank test p= 0.66). **c.** Chemo-convulsant challenge with loxapine (Kaplan-Meier curve, p= 0.34, Log-rank test). **d.** Susceptibility to thermogenic seizures (Kaplan-Meier curve, p= 0.88, Log-rank test). **e.** The L/L genotype is found less frequently than expected for a L/+ intercross (p<0.0001, Chi-square test; n=40 litters, 261pups, average litter size of 7 pups). **f.** The three-chamber social interaction test was used to examine sociability of the mouse model. No significant difference was found in the time spent with the intruder mouse across the genotypes (p=0.207, Kruskal-Wallis test, +/+ n=8, L/+ n=11, L/L n=16). **g.** Spatial memory was not affected in the Y maze test. No differences were found on the time exploring a novel arm in the Y-maze (p=0.261, Kruskal-Wallis test with Dunn’s post hoc analysis; +/+ n= 8; L/+ n=8; L/L n=10).

**Supplementary figure 2.**
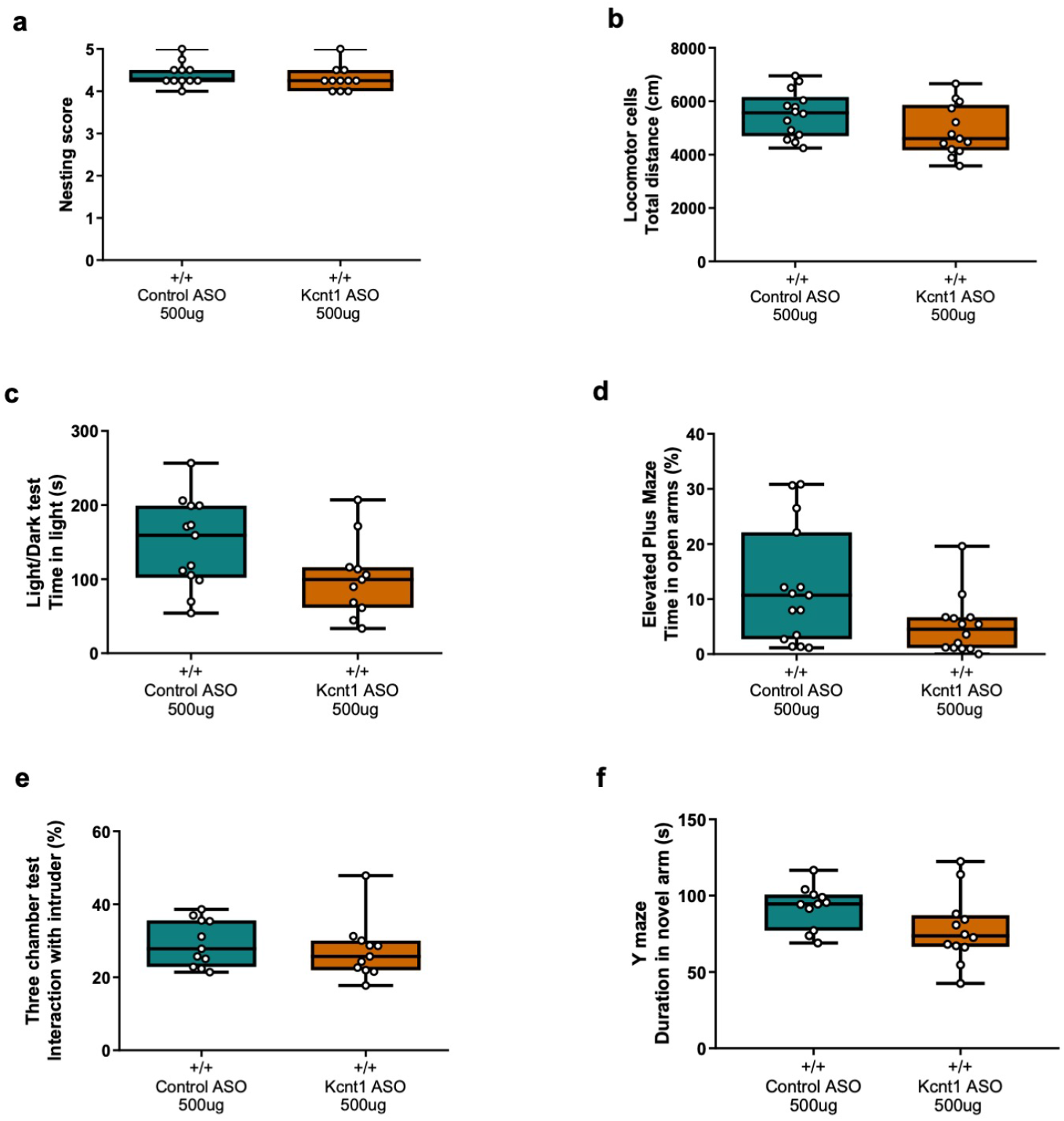
Effect of *Kcnt1* ASO administration in adult +/+ mice. **a.** Nesting behavior was not altered by *Kcnt1* ASO administration (Mann-Whitney test p=0.3136, n=11 for each group). **b.** Behavior in the locomotor cells test was not significantly affected by treatment with *Kcnt1* ASO (Mann-Whitney test p=0.0945, control ASO n=14, *Kcnt1* ASO n=13). **c.** Time spent in the light compartment was reduced although not significantly (Mann-Whitney test p=0.0821, control ASO n=13, *Kcnt1* ASO n=11). **d.** +/+ mice treated with *Kcnt1* ASO showed a reduced time in the open arms compared to control treated animals (p=0.023, Mann-Whitney test, control ASO n=15; *Kcnt1* ASO n=14). **e.** The three-chamber social interaction test showed no difference in the time spent with a stranger mouse (p=0.40,1 Mann Whitney test, control ASO n=11, *Kcnt1* ASO n=11). **f.** Time spent in the novel arm of the Y maze. +/+ mice treated with *Kcnt1* ASO spent less time in the novel arm compared to control treated mice (p=0.0439, Mann Whitney test; control ASO n=11, *Kcnt1* ASO n=12).

